# High-throughput proteomics of nanogram-scale samples with Zeno SWATH DIA

**DOI:** 10.1101/2022.04.14.488299

**Authors:** Ziyue Wang, Michael Mülleder, Ihor Batruch, Anjali Chelur, Kathrin Textoris-Taube, Torsten Schwecke, Johannes Hartl, Jason Causon, Jose Castro-Perez, Vadim Demichev, Stephen Tate, Markus Ralser

## Abstract

The ability to conduct high-quality proteomic experiments in high throughput has opened new avenues in clinical research, drug discovery, and systems biology. Next to an increase in quantitative precision, recent developments in high-throughput proteomics have also gained proteomic depth, to the extent that earlier gaps between classic and high-throughput experiments have significantly narrowed. Here we introduce and benchmark Zeno SWATH, a data-independent acquisition technique that employs a linear ion trap pulsing (Zeno trap pulsing) in order to increase proteomic depth and dynamic range in proteomic experiments. Combined with the high acquisition speed, these gains in sensitivity are particularly attractive for conducting high-throughput proteomics experiments with high chromatographic flow rates and fast gradients. We demonstrate that when combined with either micro-flow- or analytical-flow-rate chromatography, Zeno SWATH increases protein identification in complex samples 5- to 10-fold when compared to current SWATH acquisition methods on the same instrument. Using 20-min micro-flow chromatography, Zeno SWATH identified > 6,000 proteins from a 62.5 ng load of human cell lysate with more than 5,000 proteins consistently quantified in triplicate injections with a median CV of 6%. Using 5-min analytical-flow-rate chromatography (800 µl/min), Zeno SWATH identified 4,907 proteins from a triplicate injection of 2 µg of a human cell lysate; or more than 3,000 proteins from 250 ng tryptic digest. Zeno SWATH hence facilitates precise proteomic experiments with small sample amounts using a fast and robust high flow-rate chromatographic method, broadening the application space that requires precise proteomic experiments on a large scale.

## Introduction

Proteomics aims to study a large number of proteins in a biological system, with the goal of understanding protein distribution, temporal dynamics, and protein concentration in response to the environment as well as between diseased and healthy biological systems [1,2]. The study of the proteome hence requires both the ability to address its large dynamic range and complexity, as well as the dynamics of biological systems over time and in relation to the genetic background. While proteomics aims to reliably identify as many proteins as possible, many of the biological and biomedical questions that are to be addressed necessitate the processing of large sample series as well. For example, a minimal drug treatment experiment of a wild-type and a single mutant cell line, with five drug concentrations, five timepoints, and triplicate measurements already produces 150 samples. An epidemiological study such as the Fenland study [3] can reach tens of thousands of samples.

In bottom-up proteomics, data-dependent acquisition (DDA) selects and fragments multiple charge ions from a full-scan mass spectrum, which are fragmented, generating tandem (MS/MS) mass spectra that can be matched to spectra in a database. While DDA ensures selectivity, it creates problems in large-scale sample series and fast proteomic measurements due to missing values. Data-independent acquisition (DIA) techniques such as SWATH acquisition [4] address this problem by generating fragment-ion spectra from all precursor ions that fall within a predefined precursor m/z range window. DIA shifted the data processing strategies from an established spectrum-centric processing model to a peptide-centric model, requiring a shift in algorithm design and assumptions [4]. Moreover, the increased sensitivity of DIA methods has facilitated applications in large-scale proteomics, including system-biology studies in various model organisms, disease states, and species [5–9], with one of the most common clinical applications being the discovery of (compound) biomarkers for disease in body fluids, particularly blood serum or plasma [5,10–13].

Throughput, the required measurement precision, and sample amount are major factors that need to be balanced when conducting large-scale proteomics measurements. Another problem are batch effects that exist within each large proteomic experiment. Conventional DIA relies on nanoflow chromatography with increased depth of single-shot proteome analysis at low sample amounts. While nano-LC workflows dig deep in proteomics [14], the low flow rate requires longer gradient time that is susceptible to technical distortions; and columns compatible with nano-LC systems require exchange after a maximum few hundred injections, which prolongs the acquisition time and is a main source of complex batch effects in larger studies. Faster gradients and higher-flow-rate proteomics workflows have been achieved with improvement of DIA data analysis tools, simplifying high-throughput proteomics with more robust chromatography and cost-effective columns [15–17], but at the cost of a higher sample amount required.

Here, we present a new acquisition technique, Zeno SWATH, that exploits a linear ion trap, the Zeno trap, in combination with SWATH acquisition. We demonstrate that Zeno SWATH increases sensitivity and expands the dynamic range coverage of identification. With its depth, Zeno SWATH is able to perform proteomics experiments that are both fast and sensitive using micro-flow as well as high-flow chromatography.

## Results

### Implementation and function principles of Zeno SWATH

In high-throughput proteomics, the required fast acquisition speed is often reached using time-of-flight (TOF) mass spectrometers. Nonetheless, in TOF instruments all fragment ions are recorded in parallel (quasi-simultaneously), where the ions spread by m/z across the distance between the exit of the collision cell and the TOF accelerator. When the subsequent pulse is observed at the accelerator, only a small proportion of the total fragment ion population is situated at the right location to enter the TOF, leading to under-detection. This disproportionately impacts lower-m/z ions, with the duty cycle of TOFs falling in the range of only 5–25%, depending on range and m/z. Because of this duty cycle deficiency, many ions (especially low-m/z ions) are not detected in low-abundance samples, resulting in TOF-MS/MS not fully realising its sensitivity. As described by Laboda et al. [18,19], a linear ion trap introduced after the collision cell (Q2), the Zeno trap, can compensate for these limitations in TOF-MS/MS (Figure 1A). When the Zeno trap is enabled, all fragment ions in the selected mass range are trapped in an axial pseudopotential well created by an additional RF (“AC”) voltage applied with the same amplitude and phase to all four rods of the trap in a required focal point. Subsequently, the release of the ions is by potential energy with timing aligned to the next pulse at the accelerator, enabling a > 90% duty cycle, thus increasing intensity 4–20 times [19] without loss of mass accuracy and resolution. Zeno trap pulsing increases the identification of low-abundance proteins, but could also be used to lower the sample amounts required.

**Figure 1.**
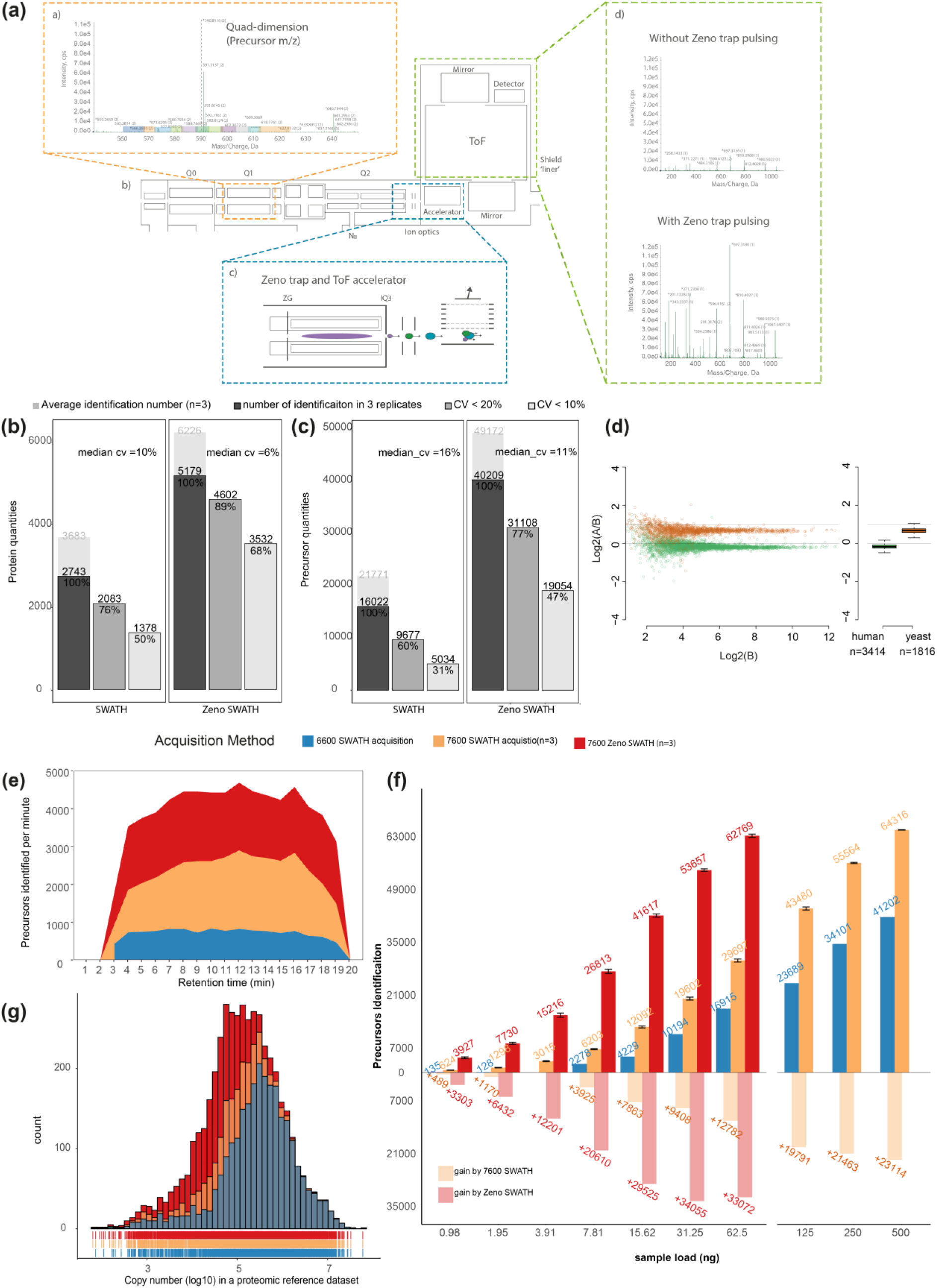
Zeno SWATH and its performance on K562 using 5 µl/min, 20 min micro-flow chromatography. **a) The Zeno SWATH DIA process.** a) Q1 quad dimension, a typical TOF-MS showing variable window distribution. b) ZenoTOF 7600 system ion path. c) Expanded view of the Zeno trap. d) MS/MS spectra for selected precursor as highlighted in a). **b**,**c) Reproducibility of protein identification using SWATH and Zeno SWATH on human cell-line standards separated by micro-flow chromatography with 62.5 ng K562 load**. Average identification numbers of proteins (b) and precursors (c) across three technical replicates (grey background bar) are given; numbers of consistent identifications in technical replicates are given in dark grey; proteins or precursors quantified with coefficient of variation better than 20% in grey, and those quantified with a CV better than 10% in light grey. **d) Protein-level LFQbench results for Zeno SWATH**. Quantification precision was benchmarked using yeast lysates that were spiked in two different proportions (A and B, three repeat injections each) into human peptide preparation (K562) (A: 30 ng K562 + 35 ng yeast; B: 30 ng K562 + 17.5 ng yeast). Raw data were processed by library-free mode DIA-NN analysis. Protein ratios between the mixtures were visualised using the LFQbench R package [21]. Left pane, log-transformed ratios (log2(A/B)) of proteins plotted for each benchmarked software tool over the log-transformed intensity of sample B. Coloured dashed lines represent the expected log2(A/B) values for human (green) and yeast (orange). Black dashed lines represent the local trend along the x axis of experimental log-transformed ratios of each population (human, yeast). Right panel, protein quantification performance shown as box plots (boxes, interquartile range; whiskers, 1–99 percentile; n = 3,414 (human) and n = 1,816 (yeast)). **e) Visualisation of precursor identification across gradient length** with SWATH and Zeno SWATH of 62.5 ng K562 injection. **f) Precursor identification performance using SWATH and Zeno SWATH**. Illustrated is the average number of precursor identifications for a K562 dilution series under three acquisition methods with library-free DIA-NN analysis. **g) Histogram of the protein abundance distribution represented as copy number in log10 scale of a human proteomic reference dataset** [24]. Zeno SWATH increases protein identification numbers by quantifying more low-abundance proteins as detected in a single-shot analysis of 62.5 ng K562 using SWATH on the TripleTOF 6600 system (blue), SWATH on the ZenoTOF 7600 system (yellow), and Zeno SWATH (red).

In order to use the Zeno trap in combination with a SWATH-MS acquisition scheme [4], we altered the acquisition software so that the Zeno trap [18] is used in a trapping and releasing setup to increase the sensitivity of MS/MS for each Q1-selected variable window acquired in a SWATH experiment (Figure 1a). Moreover, to generate platforms for medium- and high-throughput proteomic experiments, we coupled both a micro-flow LC system (ACQUITY M-Class, Waters) and an analytical-flow-rate LC (1290 Infinity II, Agilent), respectively, with a ZenoTOF 7600 system (SCIEX) equipped with a Zeno trap and the modified acquisition software. For data processing, we applied our deep-neural-network-based software suite, DIA-NN, which has been specifically optimised for handling complex data arising from fast chromatographic methods [20].

### Zeno SWATH increases proteomic depth and quantification precision

First, we examined the reproducibility of the precursor (i.e. peptide bearing a specific charge) and protein intensities between triplicate analyses of 62.5 ng of the K562 human cell line tryptic digest using Zeno SWATH. Peptides were separated using a micro-flow LC (ACQUITY M-Class, Waters), running a flow rate of 5 µl/min, 20-min micro-flow gradient chromatography (Methods). The raw data were processed with DIA-NN [20]. For analysis we used a spectral library generated using the ZenoTOF 7600 system (Methods). Quantification in Zeno SWATH mode yielded 5,179 reproducible protein IDs among an average of 6,226 proteins identified in triplicates, and reproducibility with a median coefficient of variance (CV) of 6%. Thus, 89% of the total 5,179 reproducible proteins quantified with a CV below 20%. This is an improvement over conventional SWATH, where 76% of 2,743 proteins were reproducibly quantified (CV < 20%) (Figure 1b). Indeed, the identification performance was also significantly improved. At precursor level, conventional stepped SWATH identified on average 21,771 precursors in a triplicate injection, of which 16,022 were consistently quantified in triplicates. The median CV of these precursors was 11%; 60% of the 16,022 precursors were reproducibly quantified with a CV below 20%, and 31% with a CV below 10%. In contrast, when the same injections were recorded with Zeno SWATH, on average 49,172 precursors were identified, of which 40,209 were identified consistently. Zeno SWATH also increased to 77% the percentage of precursors quantified with a CV of less than 20% (Figure 1c). These results not only demonstrate the increased sensitivity and identification rate of Zeno SWATH as compared to conventional SWATH, but also show that the increase in sensitivity simultaneously improves the quantification precision.

The analysis of different species proteomes mixed in different ratios, as implemented in the LFQbench test [21,22], has become popular in evaluating the capabilities of DIA workflows and data-processing software [20,23]. LFQbench evaluates the quantification accuracy, which is determined by the ability of the analytical setup and data processing software to deconvolute the multiplexed spectra that arise in DIA. Herein, we assessed how well the known ratios between two species lysates; human and yeast lysates mixed in the ratio 30:35 (A) and 30:17.5 (B), (Methods) were recovered in triplicates using SWATH and Zeno SWATH, followed by DIA-NN library-free analysis. A total of 3,282 human proteins and 1,692 yeast proteins were identified with Zeno SWATH, and showed valid ratios with at least two replicates in each of A and B, while only 1,741 human and 776 yeast proteins were valid in SWATH acquisition. Zeno SWATH also yielded significantly better quantification precision for human proteins compared to SWATH (Zeno SWATH, Figure 1d; SWATH, Supplementary Figure 1a): CV values for human proteins measured in samples A and B were 6.9% and 7%, respectively, using Zeno SWATH, compared to 12.7% and 12.6%, respectively, using SWATH.

### Zeno SWATH increases the acquisition sensitivity at a nanogram level of injection

Next, we tested the sensitivity of Zeno SWATH, analysing a tryptic digest of a human cell line (K562) using both SWATH and Zeno SWATH on the ZenoTOF 7600 system. K562 dilution series were acquired with a 20-min, 5 µl/min chromatographic gradient (Methods), covering sample amounts ranging from 500 ng down to near 1 ng (details shown in Methods). For a comparison of MS systems, a similar dilution series was acquired using SWATH-MS on a TripleTOF 6600 system (SCIEX) with a similar 5 µl/min, 20-min chromatography setup (Methods). To eliminate a potential bias that system-specific experimental spectral libraries have on data from different MS systems, the benchmark was carried out using library-free data analysis. At a 62.5 ng load of K562, we observed that Zeno SWATH coupled to library-free DIA-NN analysis achieves identification rates of up to 4,636 precursors detected per minute through the gradient time, whereas SWATH from the ZenoTOF 7600 system reached 2,874 precursors/min and SWATH from the TripleTOF 6600 system reached 1,153 precursors/min (Figure 1e).

The consistent gain of Zeno SWATH over SWATH acquisition in precursor identification over the 20-min active gradient was also reflected in the total number of precursor identifications. From the 62.5 ng K562, the workflow identified 16,915 precursors or 2,886 proteins using the TripleTOF 6600 system at a 1% FDR rate (Figure 1f). Using SWATH acquisition on the 7600 instrument, these numbers increased to 29,697 precursors identified from three technical replicates and on average 3,724 proteins (Figure 1f; protein quantification in Supplementary Figure 1b). With Zeno SWATH, this number increased to an average of 49,369 precursors, or 5,223 proteins, from three replicates at 62.5 ng K562 injection. Interpreting the data in a different way, Zeno SWATH identified about the same amount of precursors from 1/10 of the injected material. For instance, from 62.5 ng K562, Zeno SWATH identified 62,769 precursors or 5,206 proteins; SWATH instead identified 64,316 precursors or 5,242 proteins from 500 ng of the same sample (Figure 1f).

We speculated the identification gains could be explained by a better quantification of low-abundance proteins. We therefore assessed the dynamic range of the proteins covered by comparing our K562 results to a proteomic reference dataset with copy-number estimates for 14,178 human protein groups [24]. Conventional SWATH in the 7600 system showed a larger protein dynamic-range coverage compared to the TripleTOF 6600 system. Zeno SWATH further increased the concentration range, so that more low-abundance proteins were quantified (Figure 1g).

In summary, these results show that in general the ZenoTOF 7600 system has higher sensitivity compared to the TripleTOF 6600 system, but a higher relative gain in protein identification rates, and a higher number of precisely quantified precursors are achieved with Zeno SWATH. The gain in identification number of Zeno SWATH versus SWATH is mostly explained by an increased dynamic range: i.e. more low-abundance proteins are detected.

### Zeno SWATH facilitates high-throughput analytical-flow-rate proteomics with low sample amounts

Because of its high analytical performance and stability in combination with short chromatographic gradients, analytical-flow-rate chromatography is a preferred method in routine laboratories as well as large-scale projects. Specifically for applications such as neat-plasma proteomics or drug screens, we have previously demonstrated proteomic experiments using flow rates of 800 µl/min, and obtained quantification of precise proteomes with active gradients that ranged from 30 seconds to 5 minutes [15,25]. High-flow chromatography is attractive for practical reasons, because columns equilibrate faster, and the problem of dead volumes and sample carryover is substantially reduced. In our recent studies we were able to reduce the time between injections to 3 minutes or less [25]. As a consequence, even with the slowest of these gradients—5 minutes of active separation—the throughput of a proteomics study could largely increase to 180 samples per day in a single instrument. The downside so far is a reliance on relatively large sample amounts, which restricts the application space and shortens the instrument cleaning cycles.

We therefore examined the performance of Zeno SWATH using an analytical-flow-rate HPLC (1290 Infinity II, Agilent), running at 800 µl/min, with a 5-min gradient chromatography. We analysed K562 dilution series (2 µg down to 4 ng) raw data using spectral library-based DIA-NN. On average Zeno SWATH identified 29,681 precursors and 4,907 proteins with a 2 µg load on the column, whereas conventional SWATH identified 17,212 precursors or 3,250 proteins with the same amount of sample (protein data shown in Figure 2a, respective precursor data in Supplementary Figure 2a). The high stability of the analytical-flow-rate LC system was reflected in the reproducibility of the analysis, and achieved a median protein quantification CV of 5%. Expressed differently, 92% of the proteins were consistently identified with a CV < 20% (Figure 2b, respective precursor data in Supplementary Figure 2b). Importantly, with Zeno SWATH, analytical-flow-rate chromatography was able to process low sample amounts. For instance, with just 250 ng of digest injected, more than 3,000 proteins were quantified (Figure 2a). Thus, with Zeno SWATH, analytical-flow-rate proteomics becomes applicable for proteomic experiments that produce low amounts of samples; and when higher sample amounts are available, ID numbers close to those in conventional proteomics (low-flow-rate, long-gradient chromatography) are reached.

**Figure 2.**
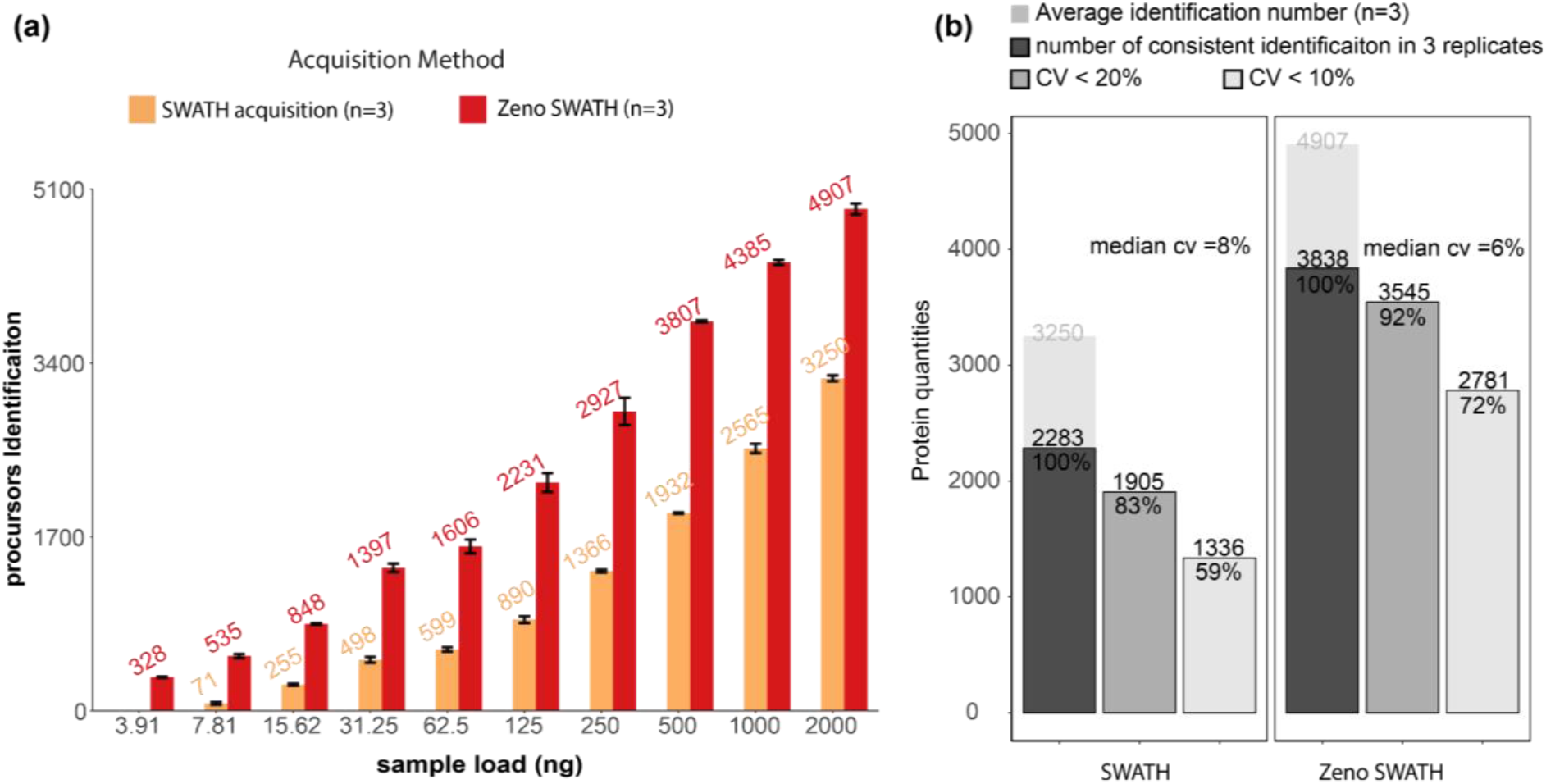
Protein identification performance of K562 using 800 µl/min, 5-min gradient chromatography in SWATH, and Zeno SWATH. **a)** Comparison of sample injection amount and identification performance with SWATH and Zeno SWATH. **b)** Reproducibility of SWATH and Zeno SWATH on K562 separated by analytical-flow-rate chromatography. average identification number in three replicates (background), number of identifications in all replicates (dark grey), proteins with coefficient of variation below 20% (grey) and below 10% (light grey) of SWATH and Zeno SWATH. Raw data were analysed by spectral library-based DIA-NN analysis.

### Evaluation of Zeno SWATH on plasma, plant, fungal, and bacterial samples

A key application space for high-throughput proteomics is human plasma and serum proteomics, as well as systems and synthetic biology. These matrices differ in dynamic range and proteomic complexity from a human cell-line extract such as K562. For instance, in human plasma, 99.9% of the proteomic mass is attributable to less than 200 proteins [26]. To assess the performance of Zeno SWATH on various matrices, we hence assayed a dilution series (1∼500 ng) of human plasma as well as cell extracts from the yeast *Saccharomyces cerevisiae*, the bacterium *Escherichia coli*, and seedlings of chickpea (*Cicer arietinum*). The standards and samples generated were acquired using the 20-min micro-flow gradient (Methods) and MS method with both Zeno SWATH and SWATH.

In non-depleted human plasma, Zeno SWATH identified 3,957 precursors derived from 204 proteins with 62.5 ng injection, covering a larger dynamic concentration range of protein than the same sample load acquired by SWATH (3,947 precursors from 245 proteins of Zeno SWATH and SWATH 2,516 precursors from 156 proteins). 15 ng of sample was already sufficient to quantify the dominating fraction of the plasma proteome (160 proteins) (Figure 3). Zeno SWATH could hence be applied for high-throughput plasma samples where sample amounts are limited, for instance to make proteomics applicable for analysing finger-prick samples or in combination with nano-sampling devices.

**Figure 3.**
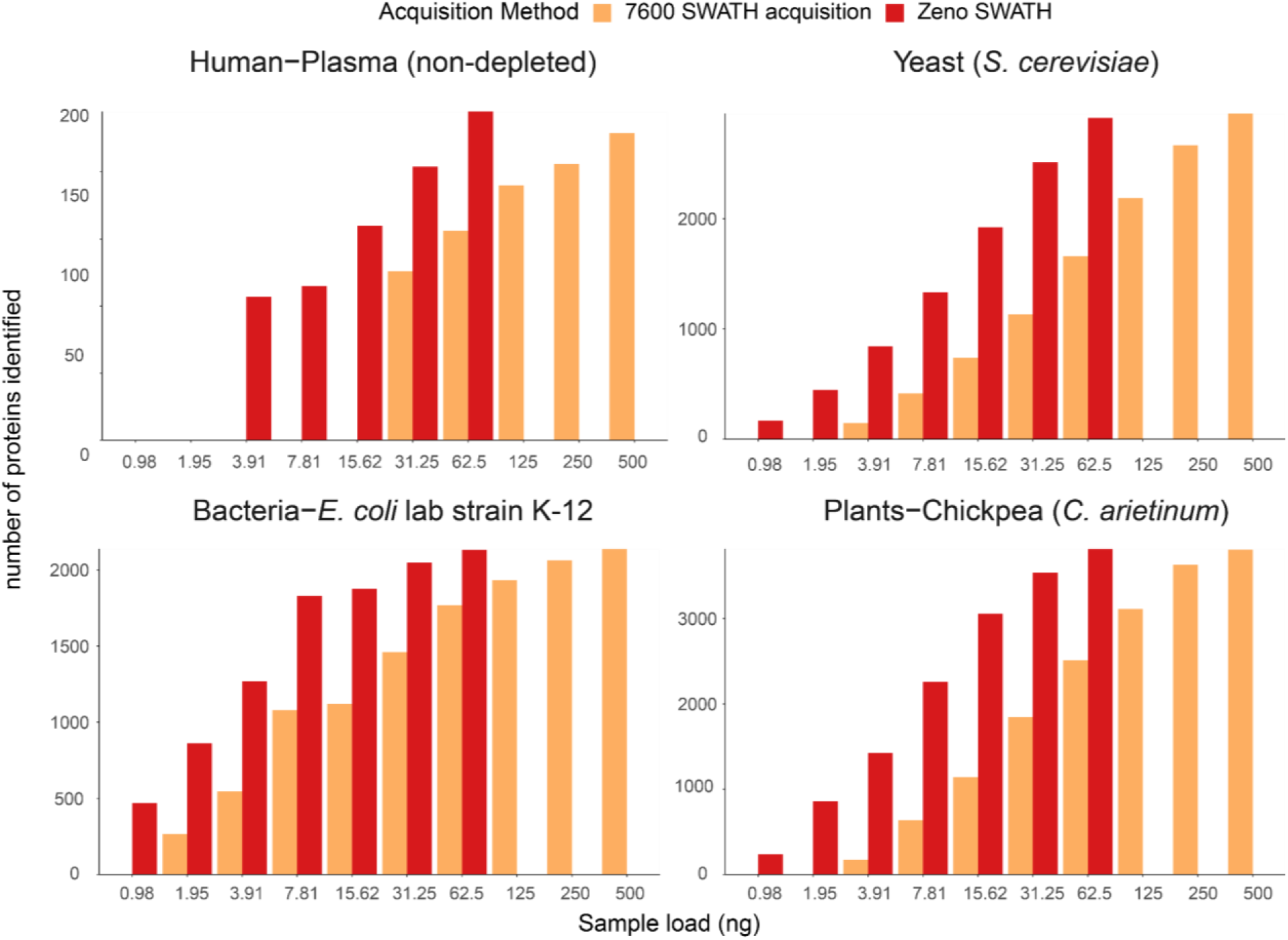
Protein identification with SWATH acquisition and Zeno SWATH in different sample types with 5-µl/min, 20-min micro-flow chromatography. We generated tryptic digests from human plasma, and protein extracts from the yeast *S. cerevisiae*, the *E. coli* lab strain K-12, and chickpea (*C. arietinum*) seedlings germinated in the lab. Data was processed with library-free DIA-NN analysis.

In *S. cerevisiae* protein extract, micro-flow Zeno SWATH quantified 29,339 precursors from 2,915 proteins from the 62.5 ng injection, compared to 1,660 proteins quantified by SWATH on the same instrument setup. In *E. coli*, Zeno SWATH identified 24,880 precursors or 2,131 proteins at 62.5 ng load, whereas conventional SWATH acquired 16,139 and 1,768 precursors and proteins, respectivtly. And finally, we identified 33,694 precursors and 3,815 proteins from 62.5 ng of *C. arietinum* samples, whereas in SWATH acquisition 500 ng of sample was needed to reach similar identification numbers (34,573 precursors, 3,807 proteins from SWATH) (An overview of achieved protein identification is shown in Figure 3; details for respective precursor identification are shown in Supplementary Figure 3; identification numbers for both proteins and precursors are shown in Supplementary Table 1). Hence, despite the varying complexity and differences in the dynamic range of the different species proteomes, Zeno SWATH largely increased protein identification rates, or enabled similar identification numbers compared to SWATH using much lower sample amounts.

### Conclusions

Proteomic experiments are regularly limited by sample amount or instrument sensitivity. We and others have previously presented high-throughput proteomic experiments using micro-flow-[16,17] and analytical-flow-rate chromatography [15,25]. Specifically the latter presents highly desirable chromatographic properties, but requires high sample amounts. We therefore sought for new ways to maintain the performance of the high-throughput proteomic strategies, but to decrease their reliance on high sample amounts, or alternatively, increase sensitivity and proteomic depth. Here, we presented a Zeno-trap-enabled DIA acquisition strategy (“Zeno SWATH”) implemented on the ZenoTOF 7600 system as a way to increase protein identification in fast proteomic experiments. Testing Zeno SWATH on a digested human protein extract, on human plasma, as well as on yeast, *E. coli*, and chickpea seedling samples, we report that, on average, Zeno SWATH quantifies a similar number of precursors from about 1/10 of the injection amount in medium- and high throughput experiments compared to SWATH-MS. Protein identification numbers, as well as the number of precisely quantified peptides, increase significantly when the same amount of sample is injected in Zeno SWATH. Zeno SWATH facilitates proteomic experiments in conjunction with analytical-flow-rate chromatography, where so far sample amounts required were limiting and identification rates lower.

In summary, Zeno SWATH can improve proteomic experiments in several ways. It increases the number of protein identifications as well as the number of precisely quantified peptides with no extra effort when compared to a SWATH experiment, using micro-flow chromatography. Furthermore, in large-scale proteomic studies, Zeno SWATH can be used in high-throughput proteomic experiments, reducing the sample-amount requirements and increasing protein identification numbers through the use of analytical-flow-rate chromatography.

Further improvements in sensitivity in high-throughput proteomics are still possible in the future. For instance, our lab has previously introduced a Scanning SWATH acquisition technique that significantly increased the identification rate of proteins using the TripleTOF 6600 system (SCIEX), particularly in ultra-fast proteomic experiments [25]. Scanning SWATH combines the benefits of data-dependent and data-independent techniques by employing a ‘sliding’ quadrupole (Q1) to assign precursor masses to MS/MS traces, allowing for a faster Q1 scan than stepwise SWATH acquisition. A combination of the Zeno trap trapping technology with Scanning SWATH is technically possible, and might result in further gains in the sensitive analysis of proteomes in high throughput.

## Acknowledgements

We thank the Charité Core Facility High Throughput Mass Spectrometry for technical support, and advice in experimental design, sample preparation, data analysis, and interpretation; We thank Fatma Amari, Spyros Vernardis and Poppy Adkin for data acquisition from TripleTOF 6600 system; Lisa Juliane Kahl and Sreejith J. Varma for providing *E*.*coli* and *C. arietinum* seedling stocks; and Hezi Tenenboim as well as our team members for proofreading and commenting on the manuscript.

This work was supported by the Ministry of Education and Research (BMBF), as part of the National Research Node ‘Mass spectrometry in Systems Medicine’ (MSCoreSys), under grant agreements 031L0220 (to M.R.) and 161L0221 (to V.D.), and the Deutsche Forschergemeinschaft (DFG) as part of the Sonderforschungsbereich (SFB) TRR-185. ZW is a member of the International Max Planck Research School (IMPRS) for Infectious Diseases and Immunology. Work in the MR lab is further supported by the Francis Crick Institute, which receives its core funding from Cancer Research UK (FC001134), the UK Medical Research Council (FC001134), the Wellcome Trust (IA 200829/Z/16/Z) and the European Research Council (ERC-SyG-2020 951475).

## Contributions

Conceptualisation & Supervision: MM, VD, MR ;Data curation & Formal analysis: ZW; Funding acquisition: VD, MR; Investigation: ZW, KTT (data acquisition); TS (sample preparation); Methodology: ZW, KTT, IB; Project administration: MM, JC, JCP, ST, MR; Resources: IB, AC, JC, JCP, ST; Software: VD; Visualisation: ZW, MM, JH; Writing – original draft, editing, proofreading: ZW, TS, JH, JC, VD, MR

## Materials and methods

### Materials and reagents

Water was from Merck (LiChrosolv LC-MS grade; Cat# 115333), acetonitrile was from Biosolve (LC-MS grade; Cat# 012078), 1,4-Dithiothreitol (DTT; Cat# 6908.2) from Carl-Roth, iodoacetamide (IAA; Bioultra; Cat# I1149) and urea (puriss. P.a., reag. Ph. Eur.; Cat# 33247) were from Sigma-Aldrich, ammonium bicarbonate (Eluent additive for LC-MS; Cat# 40867), formic acid (LC-MS Grade; Eluent additive for LC-MS; Cat# 85178) was from Thermo Fisher Scientific, trypsin (sequence grade; Cat# V511X), digested human protein extract (K562, Mass Spec-Compatible Human Protein Extracts, Cat# V6951) and digested yeast protein extract (Mass Spec-Compatible Yeast Protein Extracts, Cat# V7461) were from Promega, commercial human plasma samples (Human Source Plasma, LOT# 20CILP1034) was from ZenBio.

### Sample preparation

**K562 and yeast** stock samples were prepared via reconstitution of the powder form of the product using 200 µl of reconstitution buffer (1% ACN, 0.1% formic acid in LC grade Water) and aliquoted into 4 × 50 µl stock sample, stored at –80°C and defrosted before sample preparation.

**Plasma** stock samples were prepared as follows: 5 µl of commercially available plasma sample was added to 55 µl of denaturation buffer (50 µl of 8 M urea, 100 mM ammonium bicarbonate (ABC), 5 µl of 50 mM dithiothreitol (DTT)). The samples were incubated for 1 h at room temperature before the addition of 5 µl of 100 mM iodoacetamide (IAA). After a 30 min incubation at room temperature in the dark, the samples were diluted with 340 µl of 100 mM ammonium bicarbonate and digested overnight with 22.5 µl of 0.1 µg/µl trypsin at 37°C. The digestion was quenched by adding 50 µl of 10% formic acid. The resulting tryptic peptides were purified on a 96-well C18-based solid-phase extraction (SPE) plate (BioPureSPE Macro 96-well, 100 mg PROTO C18, The Nest Group). The purified samples were resuspended in 120 µl of 0.1% formic acid.

***E. coli* samples** were prepared from 10^10^ cells, thawed in 200 µl lysis buffer (7 M urea, 0.1 M ABC, 2 mM MgCl2, 0.125 U benzonase). The cell pellets were disrupted using a GenoGrinder in the presence of 100 mg of 0.1 mm ceramic beads. The lysate was incubated at 37°C for 15 min to ensure optimal benzonase activity. The supernatant was clarified by centrifugation at 4°C, 20,000 *g* for 15 min.

Following addition of DTT to a final concentration of 5 mM the samples were incubated for 90 min at room temperature before the addition of IAA to a final concentration of 10 mM. After 30 min incubation at room temperature in the dark, the samples were diluted 5-fold with 100 mM ABC and digested overnight with 0.1 µg/µl trypsin at 37°C. The digestion was quenched by adding formic acid to a final concentration of 1%. The resulting tryptic peptides were purified using C18-based (AttractSPE, Affinisep) stage tips as described by Rappsilber et al. [27], and dried before resuspension in 0.1% formic acid.

**Chickpea seedling sample preparation** was performed essentially as described by Wang et al. [28]. Briefly, 500 mg of chickpea seedlings, created by the germination of chickpeas in the lab, were pulverised in liquid nitrogen. The powder was washed with trichloroacetic acid/acetone, methanol, and acetone at 4°C. After removal of acetone, the pellet was dried at 50°C for 20 min, the protein was subjected to phenol extraction and precipitated by the addition of ice-cold 0.1 M ammonium acetate in 80% methanol. The resulting pellet was washed once with ice-cold methanol and once with ice-cold 80% acetone, then air-dried.

The protein pellet was dissolved in 8 M urea, 100 mM Tris, pH 8.0. Following dilution to 2 M urea, the protein was reduced with 2 mM dithiothreitol (DTT) for 30 min at room temperature and alkylated for 30 min with 15 mM chloroacetamide. The sample was then digested overnight with LysC and trypsin. The resulting peptides were filtered through a 30-kDa MWCO centrifugal filter and further purified by C18 stage tips as described above.

**LFQbench hybrid samples**: to generate the hybridisation of different species samples, tryptic peptides were combined in the following ratios: MIX A was composed of 30 ng/µl of K562 and 35 ng/µl yeast tryptic digest; MIX B was composed of 30 ng/µl of K562 and 17.5 ng/µl yeast tryptic digests.

#### Instrumentation

### Micro-flow system

K562 dilution series and all sample species were acquired on an ACQUITY UPLC M-Class system (Waters) coupled to a ZenoTOF 7600 mass spectrometer with an Optiflow source (SCIEX). Prior to MS analysis, samples were chromatographically separated with a 20-minute gradient ((time, % of mobile phase B: 0 min, 3%; 0.86 min, 7.1%; 2.42 min, 11.2%; 5.53 min 15.3%; 9.38 min, 19.4%; 13.02 min, 23.6%; 15.48 min, 27.7%;17.27 min, 31.8%; 19 min, 40%; 20min, 80%), Supplementary table 2) on HSS T3 column (300 µm × 150 mm, 1.8 µm, Waters) heated to 35°C, using a flow rate of 5 µl/min [8] where mobile phases A and B are 0.1% formic acid in water and 0.1% formic acid in ACN, respectively. To compensate for injection volume variance, 1 µl of each dilution series sample or mixture sample was loaded prior to samples entering the MS. A SWATH acquisition scheme with 60 variable size windows covering a precursor mass range of 400–900 m/z (Supplementary Table 2) and 11 ms accumulation time was used. A Zeno SWATH acquisition scheme with 85 variable size windows and 11 ms accumulation time was used (Supplementary Table 1). For both methods, ion source gas 1 and 2 were set to 12 and 60 psi, respectively; curtain gas to 25, CAD gas to 7, and source temperature to 150°C; spray voltage was set to 4,500 V. K562 dilution series were acquired on a nanoAcquity UPLC System (Waters) coupled to a SCIEX TripleTOF 6600 mass spectrometer. The peptides were separated with the same 20-min gradient on a Waters HSS T3 column (300 µm × 150 mm, 1.8 µm) using a flow rate of 5 µl/min. SWATH MS/MS acquisition scheme with 40 variable size windows and 35 ms accumulation time was used [8].

### High-flow system

Standards and samples were acquired on an 1290 Infinity II UHPLC system (Agilent) coupled to a ZenoTOF 7600 mass spectrometer with a DuoSpray TurboV source (SCIEX). Prior to MS analysis, samples were chromatographically separated on an Agilent InfinityLab Poroshell 120 EC-C18 1.9 µm, 2.1 mm × 50 mm column heated to 30°C. A 5-min 0.8 ml/min flow rate of linear-gradient ramping from 3% ACN/0.1% formic acid to 40% ACN/0.1% formic acid was applied [25]. 1 µl of each dilution series sample was loaded prior to samples entering MS. A SWATH acquisition scheme with 60 variable size windows and 11 ms accumulation time was used (Supplementary Table 1). The Zeno SWATH scheme has the same 60 windows as the SWATH method and 13 ms accumulation time. For both acquisition methods, ion source gas 1 (nebuliser gas), ion source gas 2 (heater gas), and curtain gas were set to 60, 65, and 55 psi, respectively; CAD gas was set to 7, source temperature to 700°C, and spray voltage to 3,500 V.

#### Spectral libraries

The library for the K562 benchmark experiments was generated from in-house pH fractionated HeLa (Thermo Fisher Scientific, #88329) and K562 cell lines. Individual pH fractions were analysed with ZenoTOF 7600 in IDA/DDA mode and searched with OneOmics against a human Swiss-Prot canonical and isoform FASTA file (July 2020).

For the analysis of plasma samples, a project-independent public spectral library [29] was used as described previously [15]. The Human UniProt [30] isoform sequence database (UP000005640, 19 October 2021) was used to annotate the library and the processing was performed using the MBR mode in DIA-NN. For the analysis of samples from other species, the respective FASTA from Uniprot (yeast: UP000002311_559292 (28 September 2021) *E. coli*: UP000000625 (19 October 2021); chickpea: UP000087171_3827 (22 October 2021)) was used to analyse data in library-free mode.

For the analysis of hybrid species samples, in silico-predicted combined spectral library out of a combination of human and yeast FASTA were created with the enable of the deep-learning-based spectra, RTs and IMs prediction option in the Precursor ion generation pane.

#### Data processing and analysis

All raw data were processed with DIA-NN (v1.8 beta 20 provided in the Supplementary Materials) [20] using the default settings with a fragment ion m/z range of 100–1800, mass accuracy of 20 and 12 ppm at the MS2 and MS1 level, respectively, scan window of 7, protein inference disabled (spectral library) or relaxed (library free), and quantification strategy of “robust LC (high precision)". K562 acquisitions were analysed both using a spectral library and library-free mode, while all other runs were processed in the library-free mode.

Coefficients of variation (CV) were calculated for each protein or precursor as its empirical standard deviation divided by its empirical mean, and are reported in percentages. CV values were calculated for proteins or precursors identified in all 3 replicate measurements.

