## Supplementary Figures for "High-throughput proteomics of nanogram-scale samples with Zeno SWATH DIA"

Supplementary Figure 1. Zeno SWATH acquisition and its performance on K562 using 5µl/min, 20-min micro-flow chromatography.

Supplementary Figure 2. Precursors identification performance of K562 at 800 µl/min, 5-min gradient chromatography in SWATH acquisition, and Zeno SWATH.

Supplementary Figure 3. Precursors identification with SWATH acquisition and Zeno SWATH in different sample types with 5µl/min, 20-min microflow chromatography.

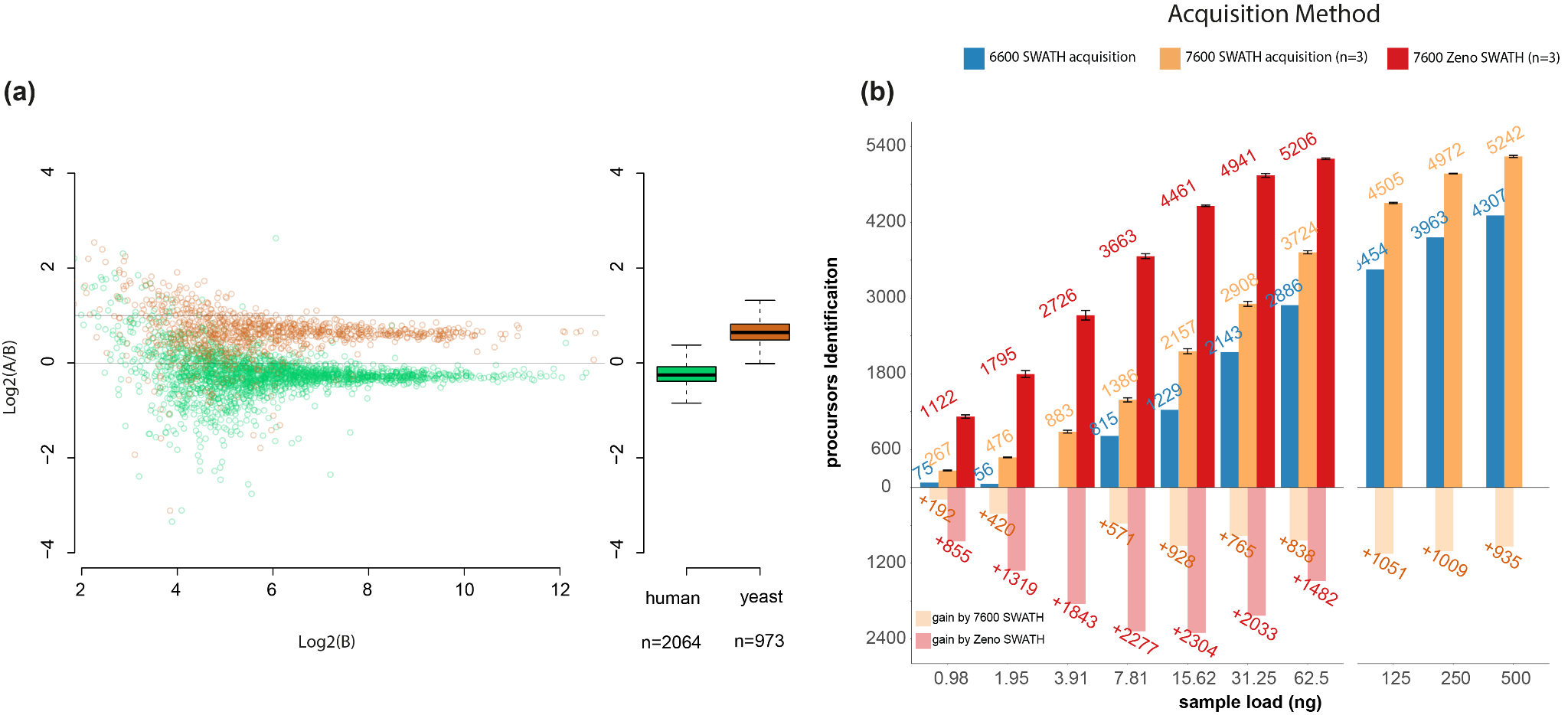

**Supplementary Figure 1. Zeno SWATH acquisition and its performance on K562 using 5µl/min, 20-min micro-flow chromatography.** **a) Protein-level LFQbench results for SWATH acquisition.** Quantification precision was benchmarked using yeast lysate that were spiked in two different proportions (A and B, three repeat injections each) into a human peptide preparation (A: 30 ng K562 + 35 ng yeast; B: 30 ng K562+ 17.5 ng yeast). Raw data were processed by library-free mode DIA-NN analysis. Protein ratios between the mixtures were visualised using the LFQbench R package [[22]](https://paperpile.com/c/IaZEo2/vFAl). Left pane, log-transformed ratios (log2(A/B)) of proteins plotted for each benchmarked software tool over the log-transformed intensity of sample B. Colored dashed lines represent the expected log2(A/B) values for human (green), yeast (orange). Black dashed lines represent the local trend along the x axis of experimental log-transformed ratios of each population (human, yeast). Right panel, protein quantification performance shown as box plots (boxes, interquartile range; whiskers, 1–99 percentile; n = 2,064(human) and n = 973(Yeast)). **b) Protein identification performance using SWATH and Zeno SWATH.** Illustrated is the average number of precursors identifications for K562 dilution series under 3 acquisition methods with library-free DIA-NN analysis.

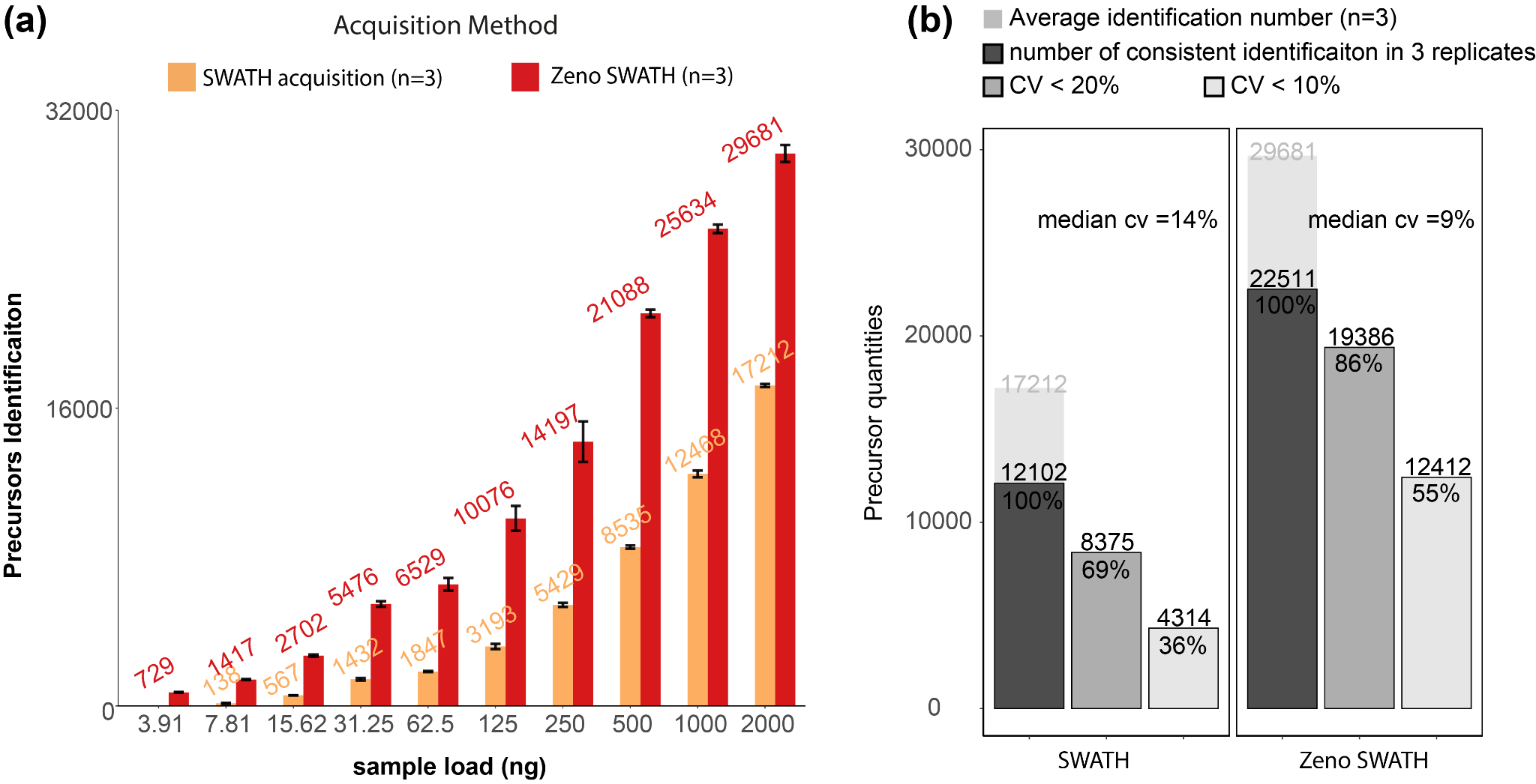

**Supplementary Figure 2. Precursors identification performance of K562 at 800 µl/min, 5-min gradient chromatography in SWATH acquisition, and Zeno SWATH. a)** Dependency of sample injection amount and identification performance on SWATH acquisition and Zeno SWATH. **b)** Reproducibility of SWATH and Zeno SWATH on K562 separated by high flow chromatography. Bar chart of average identification number in 3 replicates (background), number of identification in all replicates (dark grey), proteins with coefficient of variation below 20% (grey) and below 10% (light grey) of SWATH acquisition and Zeno SWATH. Raw data were analysed by spectral library-based DIA-NN analysis.

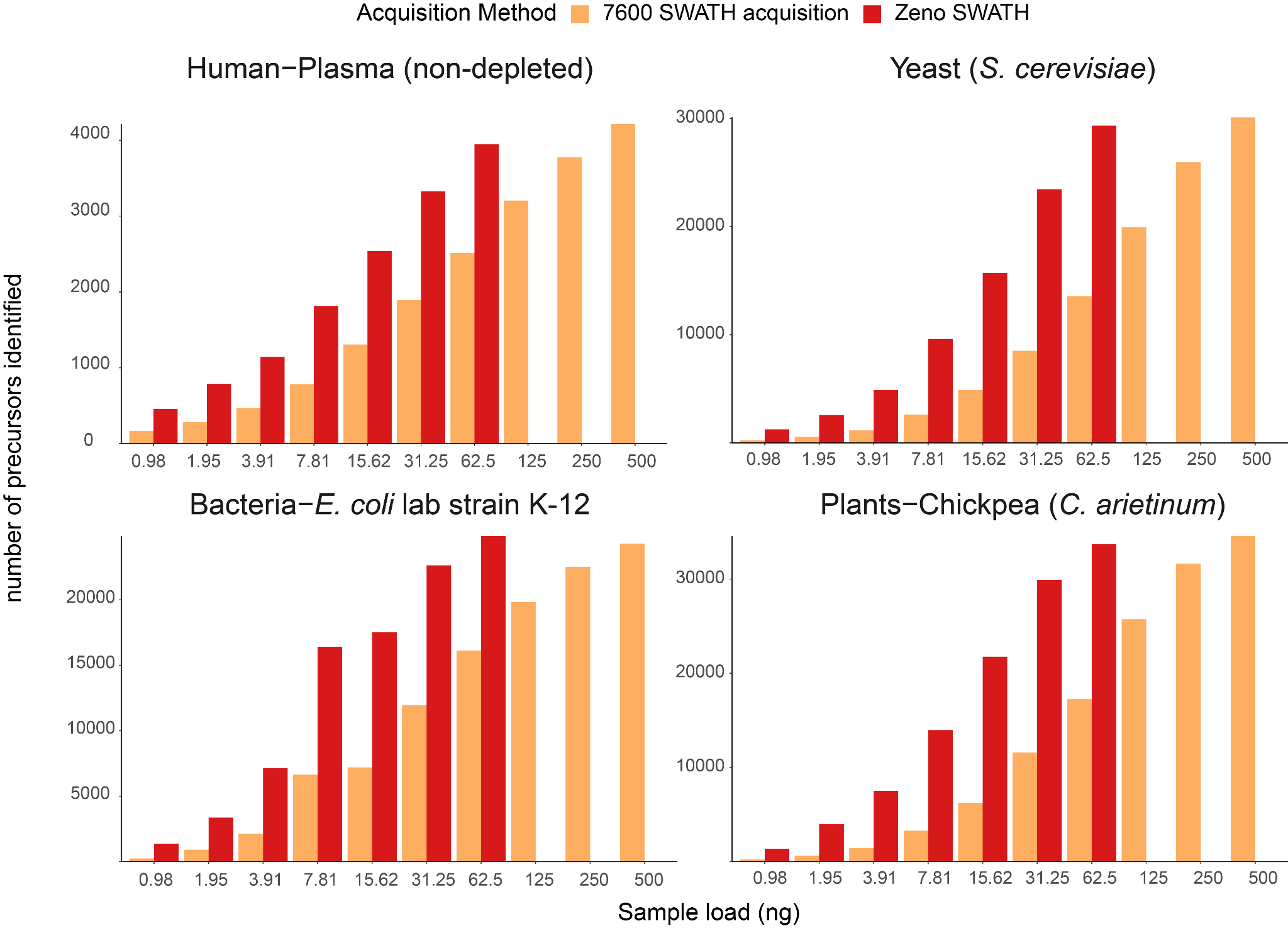

**Supplementary Figure 3.** **Precursors identification with SWATH acquisition and Zeno SWATH in different sample types with 20-min microflow chromatography.**  We generated tryptic digests from human plasma, and protein extracts from the yeast *S. cerevisiae*, the *E. coli* lab strain K-12, and Chickpea (*C. arietinum*) seedling germinated in the lab. Illustrated are the proteins identified using SWATH acquisition and Zeno SWATH with 20-minute microflow chromatography. Data was processed with library-free DIA-NN analysis.
